# Resistance genes in global crop breeding networks

**DOI:** 10.1101/106484

**Authors:** K. A. Garrett, K. F. Andersen, F. Asche, R. L. Bowden, G. A. Forbes, P. A. Kulakow, B. Zhou

**Affiliations:** First and second authors: Plant Pathology Department, University of Florida, Gainesville, FL 32611-0680, United States; First, second, and third authors: Institute for Sustainable Food Systems, University of Florida, Gainesville, FL 32611-0680, United States; First and second authors: Emerging Pathogens Institute, University of Florida, Gainesville, FL 32611-0680, United States; Third author: School of Forest Resources and Conservation, University of Florida, Gainesville, FL 32611, United States; Fourth author: USDA-ARS Hard Winter Wheat Genetics Research Unit, 4008 Throckmorton Hall, Kansas State University, Manhattan, KS 66506, United States; Fifth author: International Potato Center (CIP), Kunming, China; Sixth author: International Institute of Tropical Agriculture (IITA), Ibadan, Nigeria; Seventh author: International Rice Research Institute (IRRI), Manila, Philippines

## Abstract

Resistance genes are a major tool for managing crop diseases. The crop breeder networks that exchange resistance genes and deploy them in varieties help to determine the global landscape of resistance and epidemics, an important system for maintaining food security. These networks function as a complex adaptive system, with associated strengths and vulnerabilities, and implications for policies to support resistance gene deployment strategies. Extensions of epidemic network analysis can be used to evaluate the multilayer agricultural networks that support and influence crop breeding networks. We evaluate the general structure of crop breeding networks for cassava, potato, rice, and wheat, which illustrate a range of public and private configurations. These systems must adapt to global change in climate and land use, the emergence of new diseases, and disruptive breeding technologies. Principles for maintaining system resilience can be applied to global resistance gene deployment. For example, both diversity and redundancy in the roles played by individual crop breeding groups (public versus private, global versus local) may support societal goals for crop production. Another principle is management of connectivity. Enhanced connectivity among crop breeders may benefit resistance gene deployment, but increase risks to the durability of resistance genes without effective policies regarding deployment.

## Introduction

Epidemiological network analysis offers an important perspective in plant pathology, and is becoming a standard tool for understanding the spread of disease (Moslonka-Lefebvre et al. 2011; Shaw and Pautasso 2014). Epidemic network analysis provides insights into not only disease spread between individual pairs of locations, but also the cumulative effects of connections between locations that influence regional processes and determine whether regional disease management is successful. Multilayer networks can be used to integrate understanding of system components, such as how the spread of information influences the spread of disease (Garrett 2012, 2017; Hernandez Nopsa et al. 2015). Another multilayer network that drives epidemics, and the potential for their successful management, is the movement of disease and pest resistance genes through the components of crop breeding networks.

This paper addresses the global crop breeding network, the global set of crop breeder groups and the links formed between them by the movement of genes in crop germplasm, which is a major factor in determining the global distribution of crop genotypes and phenotypes (Fig. 1). While gene networks within individual organisms are a growing research focus, the global crop breeding network has received limited analysis from a systems standpoint, despite its key role in food security during global change (Fowler and Hodgkin 2004). Resistance genes offer one of the most sustainable approaches to management of diseases and arthropod pests (Boyd et al. 2013; Byerlee and Dubin 2009), although resistance genes have variable lifespans, some lasting only a few seasons while others remain effective for decades (McDonald and Linde 2002). In some cases, the deployment of resistance genes can be coordinated in an effectively structured crop breeding network to decrease the likelihood that pathogen or pest populations evolve to overcome resistance. When new germplasm is introduced into a region it may bring desirable traits (increased yield, drought resistance, vigor) but may inadvertently introduce susceptibility to endemic pathogens, resulting in new disease outbreaks. Alternatives to host plant resistance for disease and pest management often have economic or environmental costs: use of foliar and seed-applied pesticides may impact non-target species and cultural practices such as increased tillage may increase soil erosion. These alternatives may also be too expensive, especially for smallholder farmers in developing countries. Host plant resistance, on the other hand, generally has little additional cost above and beyond the cost of seed.

**Figure 1.**
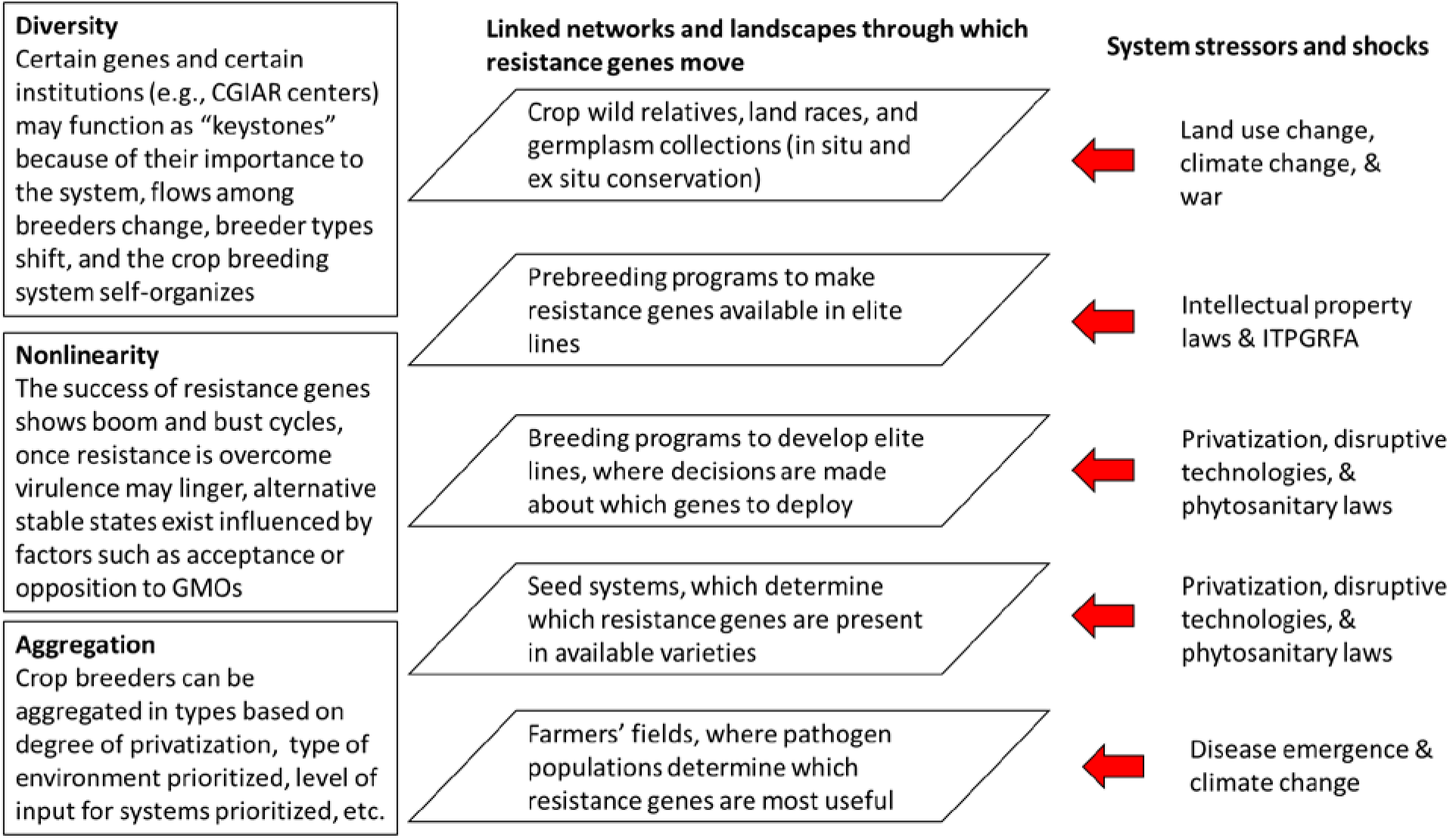
System of resistance gene deployment, including (left) characteristics of the system that make it a complex adaptive system (CAS), and (right) shocks and stressors to which the system must adapt

Several components of global change pose new challenges for the development of effective crop varieties (Fig. 1). With climate change, the geographic distribution of high risk regions for diseases and pests will shift, there will be higher uncertainty about risk influencing decision-making, and higher levels of risk than previously observed may occur (Bebber et al. 2013; Chapman et al. 2012; Garrett et al. 2013; Pautasso et al. 2012). With increasing global trade and transportation, pathogen and pest invasions will accelerate. Both land use change and climate change are likely to decrease *in situ* conservation of crop wild relatives that are a source of new resistance genes (Dempewolf et al. 2014; Jarvis et al. 2008). Crop breeding, itself, is undergoing dramatic changes. Gene editing technologies, and societal responses to them, will have major impacts on crop breeding in the near future. The steady privatization of crop breeding systems (Marden and Godfrey 2012) has important and little-studied implications for the deployment of resistance genes, and how seeds and information are shared in the crop breeding network. While public breeding groups have often had clear incentives to share resistance genes with other breeding groups, private breeding groups may have different profiles of incentives for and against sharing germplasm. Private breeding groups may also be commercially aligned with groups that develop pesticides, and in some cases overall profits to a commercial entity may be higher from investments in research and development for pesticides than for resistance genes. A lack of resistance may also incentivize farmers to buy seed more frequently to avoid the buildup of seedborne disease, providing a potential conflict of interest. As a higher proportion of crop breeders become private crop breeders, there also may be fewer crop breeders with the professional responsibility to provide a public critique of global crop breeding systems. Conversely, private crop breeding groups may be prepared to invest more in the development and deployment of resistance genes, and generally have access to larger financial resources as they can utilize capital markets if they can demonstrate expected profitability.

The science of complex adaptive systems (CAS) conceptualizes systems of agents (for example, crop breeders) who can act independently, and whose actions in aggregate produce system outcomes (Levin 1998; Miller and Page 2007). This paper draws on theory related to CAS, and the implications of such a structure for resilience strategies. The objectives of this paper are to (1) develop a framework for evaluating crop breeding networks for the spread and deployment of resistance genes, and the resulting effects on global crop epidemics; (2) evaluate a coarse representation of the global crop breeding networks for four major food crops: cassava, potato, rice, and wheat; and (3) draw on theory about system resilience to inform regional and global policies aimed at improving the deployment of resistance genes.

## Resistance genes in crop breeding networks as a complex adaptive system

Complex adaptive systems (CAS) are systems that have a set of defining traits (Fig. 1) such as hierarchies, nonlinearity, diverse identities of system agents who make choices based on their own “models” about their perceived environment, and components that can be aggregated to function differently in response to environmental stimuli (Holland 1995; Levin 1998). This set of traits may allow systems to adapt as a whole to perform aggregate functions better, though, of course, if the incentives and decision-making models that drive agents’ choices do not function well (McRoberts et al. 2011), the system as a whole may not function well, either. Puettmann et al. (2013) discuss the change in perspective from adding CAS considerations in the context of forestry: “Examples of the implications of such changes include an emphasis on multiple temporal, spatial and hierarchical scales; more explicitly considering interactions among multiple biotic and abiotic components of forests; understanding and expecting non-linear responses; and planning for greater uncertainty in future conditions.” Conceptualizing systems as CAS can facilitate analysis of system traits that are more difficult to understand from a reductionist approach (Meadows and Wright 2008).

Crop breeding networks, especially considered in terms of the multilayer networks associated with them (Fig 1), have all the traits associated with CAS. Individual crop breeding groups decide what resistance genes they will exchange with other breeding groups, and what resistance genes they will include in varieties they release. In some cases key genes will be well studied and defined, potentially with markers available for marker-assisted selection; in other cases, unknown and uncharacterized resistance genes may be exchanged and deployed, often by chance, as breeders pursue the development of high-yielding cultivars. An aggregate outcome of individual decisions about exchange of genes is the global geographic distribution of deployed major resistance genes and QTLs, and the degree to which the genes function effectively to reduce disease and support crop productivity. In addition to crop breeders, the crop breeding system includes other actors in a number of linked networks (Fig. 1). Actors involved in *in situ* and *ex situ* conservation of crop wild relatives, land races, and crop germplasm make decisions about which genes are conserved, and provide source material for pre-breeding programs. These choices are influenced by international policies, such as the International Treaty for Plant Genetic Resources for Food and Agriculture (ITPGRFA) (Esquinas-Alcázar 2005). Actors in pre-breeding programs decide which resistance genes will be included in novel genetic material available as elite breeding lines to breeding programs. Actors in breeding programs decide which genes to include in the development of commercial varieties, including which combinations of (known or unknown) genes to deploy. These decisions are generally driven by market demand for certain types of resistance, and by policies that may influence which genes are available for deployment. Agents in seed systems decide which varieties to make available in sufficient numbers for planting, and how carefully to avoid the risk of pathogen and pest spread with seed. Farmers decide which of these varieties to grow and how to manage diseases and pests (McRoberts et al. 2011; Mills et al. 2011), where demand for resistance is determined by the resulting networks of pest and disease movement (Jeger et al. 2007; Shaw and Pautasso 2014). The connectivity of resistance gene deployment in the landscape helps to determine how severe a given disease is regionally and globally, and the likelihood of pathogen or pest evolution to overcome resistance, where useful patterns of resistance deployment may provide large-scale benefits in breaking the connectivity of landscapes (Margosian et al. 2009) comparable to smaller-scale benefits from within-field cultivar mixtures (Mundt 2002; Skelsey et al. 2005). Any trait, such as drought tolerance, would have a system of feedbacks based on whether farmers perceived the varieties incorporating the trait to be good choices. Of course, epidemics have the additional feature that the good or bad disease management of neighbors will influence how well a farmer’s own management choices play out. The resulting importance of a disease to farmers provides a feedback to crop breeders in terms of their decision-making about priorities (Garrett 2017). Each actor tends to make decisions to optimize their portion of the system.

The crop breeding network as a whole helps to determine what epidemics occur and how severe they are, and the system as a whole responds to a set of challenging pulse and press stressors and shocks (Fig. 1). The multilayer network includes a range of connected components that are self-organized to a great extent, in which agents have different incentives depending on organizational structure (e.g., public vs. private, local vs. global). They adapt to new scenarios such as emerging diseases and climate change, and there are many uncertainties about the success of the systems and of different candidate strategies for gene deployment regulation. As breeding programs rely more on genomic selection and prediction of phenotype based on genotype, and less on breeder selection across a broad range of testing locations, this may result in loss of resistance that would have been naturally selected for in early screening phases (Heffner et al. 2009). In addition to the movement of genetic material, the associated movement of information about crop phenotypes, and information about the progress of epidemics or infestations, is a key system component (Deans et al. 2015; Garrett 2012).

## Methods: Modeling crop breeding networks

We collected expert perceptions about the structure of global crop breeding networks from personnel in our respective institutions with direct experience, for four major food crops: cassava, potato, rice, and wheat. These perceptions can be interpreted as representing the actual networks with coarse resolution; that is, the network representations represent the structure of global crop breeding networks in broad strokes. We focus on current forms of exchange, with the understanding that historically movement of genes was primarily from crop and pathogen centers of origin (Thormann et al. 2015). The general numbers of crop breeding groups are represented by continent, and the general tendency for links (representing the flow of genetic material) between crop breeding groups are also represented. However, individual nodes, beyond obvious hub nodes such as the CGIAR centers, are not intended to represent specific crop breeding groups. Instead, links among smaller crop breeding groups were generated randomly at representative rates in adjacency matrices plotted using the igraph package (Csárdi and Nepusz 2006) in the R programming environment (R Core Team 2016).

## Results and lessons from four crop breeding systems

To illustrate the challenges that crop breeding networks must respond to, and the general nature of current networks, we summarize the resulting structure of four crop breeding networks central to global food security (Box 1 / Fig. 2), and current challenges that these crop breeding networks must address.

**Figure 2 / in Box 1.**
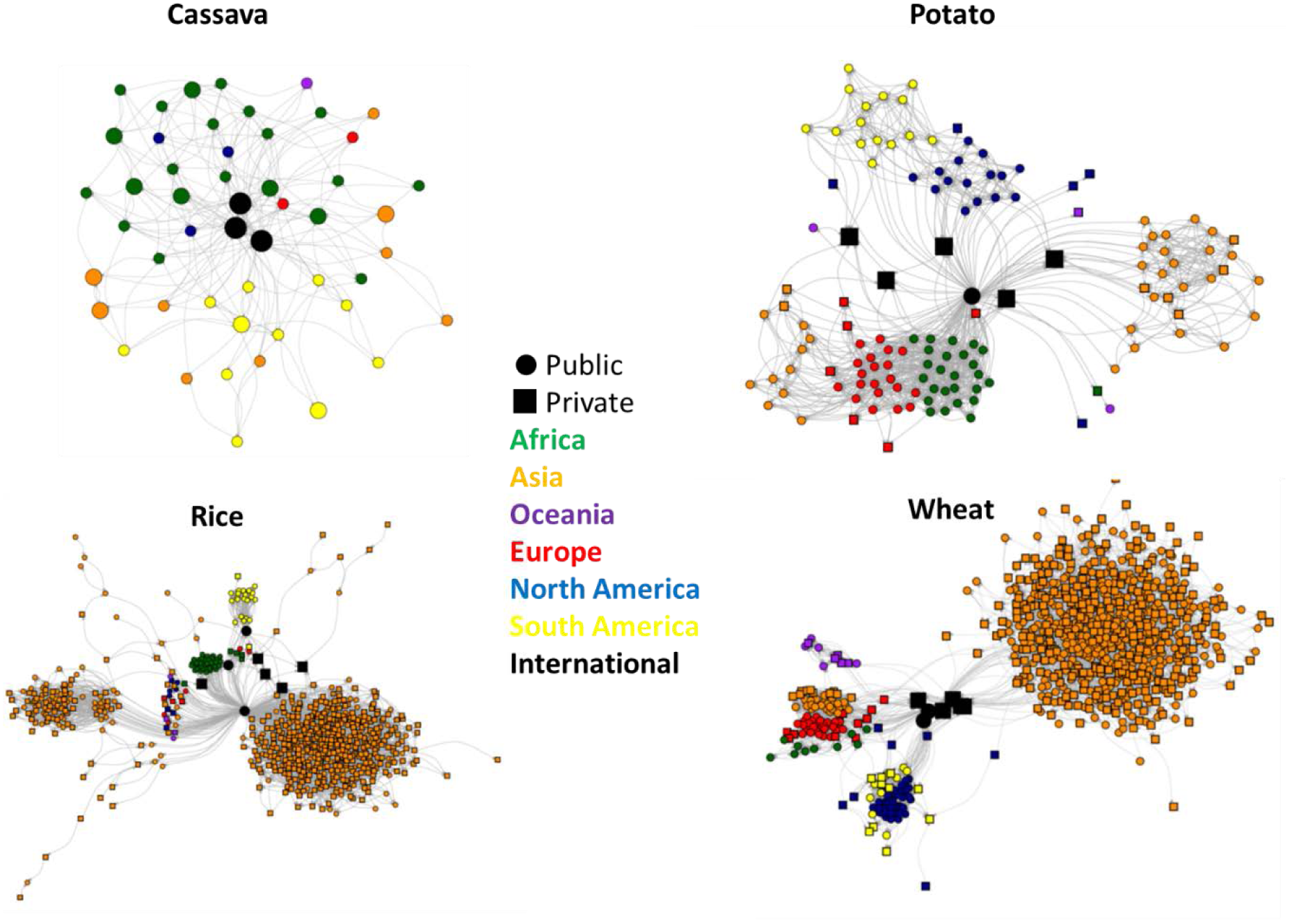
Schematics of crop breeding networks for the movement of resistance genes. These diagrams represent in coarse resolution the general structure of these networks. Each node represents a crop pre-breeding or breeding group and links represent potential dispersal of resistance genes between groups. In the **cassava** network, there are no private breeders, and a limited number of breeders, overall. In the **potato** network, both public and private potato breeders exist in high-income countries, but in low-income and middle-low income countries, there are primarily public breeding programs and few if any private potato breeders. This is because formal seed systems either do not exist or provide seed to a very small number of farmers (Thomas-Sharma et al. 2016), thus there is no mechanism for private breeders to make profits. In the **rice** network, R genes for bacterial blight and blast have been combined in high yielding backgrounds by the public sector and made available to anyone who wants them. Private breeding companies have stayed away from rice because there is not an attractive business model for seed sales. However, companies over the last 10-15 years have dramatically increased their investments in hybrid rice seed businesses. With an increase in their interests in hybrid rice, with its proven business model, companies (multinationals and smaller companies) are taking these resources and incorporating the resistance into their hybrids. If hybrids come to dominate the market, then R genes will move very quickly. However, currently uptake of hybrids is slow because of limitations on grain quality and seed costs. Uptake of R genes in inbred rice varieties is very slow, as the uptake of these varieties from the public sector can be extremely slow (e.g., it can take 25 years for a well-appreciated variety to spread to its maximum geographic distribution). Until recently, global **wheat** breeding was dominated by publicly-funded programs. Legal protections such as Plant Variety Protection (PVP) and plant patents have encouraged investments by numerous large agricultural companies. Newer technologies such as hybrid wheat, genomic selection, and transgenics offer companies many opportunities to increase market share and profit margins. One result of the pursuit of competitive advantages is a decrease in free exchange of information and genetic resources by both public and private wheat breeding programs. Discovery, pre-breeding, and dissemination of disease resistance gene resources in wheat remains primarily the domain of publicly funded institutions such as government research agencies, universities, and international agricultural research centers.

### Cassava

The network of cassava breeders across continents shows how germplasm has moved under current exchange patterns (Fig. 2). They also reflect important restrictions on germplasm movement due to phytosanitary issues, to limit the risk of the potential movement of pests and pathogens among countries. Key factors in the cassava yield gap are cassava mosaic disease (CMD), cassava brown streak disease (CBSD), and the white fly vector *(Bemisia tabaci)* of the viruses that cause them (Legg et al. 2014). The diseases are endemic to Africa with some occurrence in India; however, there is a new report of CMD in Cambodia, representing spread to a node in a new region, where current germplasm is known to be highly susceptible (Wang et al. 2015). In Africa, the biology of the viruses and the vector is more complex, with multiple viruses and strains. The spread of CMD resistance from West Africa to East Africa may also have contributed to increased occurrence of the second virus complex causing CBSD. Changes in vector populations were probably a key driver for changing viral disease patterns in East Africa, including the role of super-abundant whitefly *(Bemisia tabaci)* populations (Legg et al. 2011). The development of resistance to CMD in Africa is important but also a potential source of vulnerability. Recent genomic analyses appear to show that the genetic basis of resistance to CMD is more narrow than initially suspected. The major source of resistance appears to be durable; however, the idea of durability is also controversial in this system due to several cases that may represent the breakdown of resistance. For pests of cassava, there are important examples of both genetic resistance and biological control methods. The use of these methods has been strongly affected by networks of researchers linked with donors who support research, in addition to farmer decision-making about technology adoption. In the absence of private breeding programs, donor decision-making about long-term and short-term strategies, and about responses to system stresses, will strongly influence the system.

### Potato

Public sector potato improvement programs (PIP) began in many low-income countries in the mid to late 20th century. Their primary source of resistance genes has been the International Potato Center (CIP), which has mined native Andean potato genetic resources and also has channeled genetic resources from other advanced breeding programs. National PIP sometimes breed varieties, but often only select from those distributed by CIP. Thus CIP and national PIP can be seen as the primary decision makers about genes to be deployed.

Germplasm exchange has generally been one-way, with bi-directional information exchange assumed, but often not fully realized (Fig. 2). In some cases, germplasm has moved across borders between recipient countries, especially when regional potato networks existed in the 1990s, but inability to meet phytosanitary requirements has minimized this movement. Thus, while international property concerns may limit gene exchange in high-income countries, institutional issues can limit gene exchange in low-income countries. These limitations may be circumvented to some extent by local farmer-to-farmer exchange of seed, but this would be strongly dependent on distance among agents and other social factors. In potato, two major cases where R genes have been important are management of potato late blight (caused by *Phytophthora infestans*) and of several potato viruses, which accumulate over successive cycles of vegetative reproduction and cause yield decline, a condition referred to as degeneration (Thomas-Sharma et al. 2016). To date at least 20 R genes for resistance to *P. infestans* have been cloned (Rodewald and Trognitz 2013) and after futile efforts to use these singly, they are now being stacked to try to attain more durable resistance (Haesaert et al. 2015). To monitor pathogen evolution to these genes, a novel host plant differential set has been developed (Zhu et al. 2015). Virus genes have not been cloned but have been identified for *Potato virus Y* (PVY) and *Potato virus X* (Kopp et al. 2015), and *Potato leaf roll virus* (PLRV) (Mihovilovich et al. 2014). These genes confer extreme or hypersensitive resistance and are widely used in low-income countries where seed systems are underdeveloped.

### Rice

Two key rice diseases, rice blast (RB) and bacterial blight (BB), caused by *Magnaporthe oryzae,* and *Xanthomonas oryzae* pv. *oryzae,* respectively, pose constant threats to stable rice production. At least 25 RB and 39 BB R genes have been molecularly characterized, and most of them control race-specific resistance activated by specific pathogen avirulence (Avr) genes (Leung et al. 2015). The availability of molecular markers tightly linked and/or specific to R genes has significantly advanced the utilization of RB and BB R genes for breeding resistant rice varieties via marker assisted selection (MAS) (Ashkani et al. 2015; Rao et al. 2014). Pathogen surveillance for determining pathogen race composition using R-gene-based differential lines and diagnosis based on pathogen Avr genes has also been implemented, which is vital for efficient R-gene deployment (Dossa et al. 2015; Selisana et al. 2017). International collaborative efforts coordinated by three CGIAR centers – IRRI, CIAT, and AfricaRice – along with National Agricultural Research and Extension Services (NARES) partners, are in place for characterizing global pathogen population diversity for the smart deployment of disease resistance genes. Some broad-spectrum R genes are frequently identified as effective in several rice growing areas, e.g., the *Pi2* and *Pi9* genes for RB in Asia, Africa, and Latin America, which in turn may unwittingly narrow the diversity of the R gene pool for rice breeding programs. Due to the arms-race nature of R/Avr interactions, utilization of a limited number of R genes could drive the emergence of the same virulent pathogen races worldwide. To avoid the risk of reliance on race-specific R genes, QTLs provide an ideal alternative form of host resistance. Pyramiding of four QTLs can provide a level of resistance similar to that provided by R genes for controlling rice blast disease (Fukuoka et al. 2015). Two QTLs (*pi21* and *Pb1*) have been included in the RB resistance breeding program at AfricaRice, promoting the diversity of the R-gene pool for durable resistance in rice (Bimpong et al. 2014).

### Wheat

Wheat rusts have been an on-going focus for wheat breeding, with dramatic new system stressors in recent years (Beddow et al. 2015; Helfer 2014; Hulbert and Pumphrey 2014). Stem rust may offer the best example of the complexities of global deployment of resistance gene resources in wheat. In the 20^th^ century, stem rust was largely brought under control through near eradication of the alternate host, barberry, and through the use of genetic resistance. One of the most important resistance genes was *Sr31* (McIntosh et al. 1995), which was durable for many decades and was arguably the most widely exploited and most valuable resistance gene in wheat. Unfortunately, this gene was typically deployed singly or with other *Sr* genes that were already defeated, and little thought was given to conscious stewardship of *Sr31.* In 1999, a new race of stem rust called Ug99 was discovered in Uganda attacking wheat lines with *Sr31.* The new race was found to be highly virulent on *Sr31* as well as most known stem rust resistance genes, leaving most global wheat varieties vulnerable (Singh et al. 2015). Led by Nobel Laureate Norman Borlaug, a new global rust initiative was founded with the goal of mobilizing new genetic resistance resources to fight Ug99. Known resistance genes *Sr24* and *Sr36* were identified as being effective against Ug99, and work began on incorporating those genes into new cultivars. However, each of those genes was soon overcome by new variants of the Ug99 lineage in East Africa, most likely because they were already deployed individually in local wheat varieties. Short-term disease control has generally been favored over long-term utility of the valuable genetic resistance resources. The global wheat germplasm exchange network has been very useful and effective (Byerlee and Dubin 2009). Theoretically, combinations of disease resistance genes are much more durable than single genes. However, this is only true if deployment of the same resistance genes singly can be prevented. Preventing single deployment can be achieved through either legal means such as patents or through community cooperation in a gene stewardship plan. An alternative approach has been to identify, accumulate, and share partial or quantitative resistance genes, which individually have small effects, but are thought to be less prone to defeat by pathogens (Singh et al. 2015).

## Increased Privatization of Plant Breeding

Since the 1970s there has been an increased shift in crop breeding R&D efforts from the public to the private sector, catalyzed largely by international changes in Intellectual Property (IP) laws introducing stronger patent protections on genetic materials, along with decreased regulation around international germplasm movement, facilitating the growth of private enterprises in this sector (Frey 1996; Heisey et al. 2001; Morris et al. 2006). For many high volume crops, in industrialized markets, private companies have all but replaced the role of the public sector in finished variety release and distribution (Heisey et al. 2001). Furthermore, there has been a recent, rapid consolidation of the seed and trait industry within the private sector, resulting in larger, consolidated germplasm/IP pools (Galushko et al. 2012; Howard 2015). The involvement of the private sector in plant breeding is highly crop and region specific, with a focus on high production crops with high acreage (corn, soybean, and wheat, for example). The entrance of private companies provides a large influx of R&D and market competition into these cropping systems, with the potential to accelerate genetic gain.

The genetic material of any breeder, institution, or nation is a highly valuable asset.

When sharing genetic material among the public sector or with the private sector, public breeders must consider IP laws and licensing costs which may in some cases prove prohibitive. A survey conducted by Galusho et al. (2012) observed a decrease in the likelihood of exchanging material in a crop system that is highly privatized, such as canola, when compared to another which is highly public, such as wheat. Private breeding research programs are influenced by the needs of the market for plant disease resistance. In general, research priorities around disease resistance will mirror current grower demands in high profit regions. The acquisition of novel disease resistance traits from the public domain is, in many cases, a good return on investment for private crop breeders. Of course, resistance genes and traits do not only flow to and from the public and private sector, but also among private sector companies.

## Evaluating and interpreting the structure of networks

The topology and other features of these crop breeding networks suggest they have certain strengths and weaknesses. For example, one trait of public breeding networks (Fig. 2) is the tendency toward modularity, where modules often occur within continents as the number of links within continents is higher than the number of links among continents. Modularity may make the networks more resilient to problems such as the spread of pests through germplasm (Ash and Newth 2007), where the problem might at least be stopped before the pest leaves a module.

These networks feature key public institutions in the CGIAR system as hubs for dispersal of resistance genes (Galluzzi et al. 2016; Renkow and Byerlee 2010), along with some large national programs (Smale and Day-Rubenstein 2002). Alternatives to conventional crop breeding networks, such as organic crop breeding networks and networks for the exchange of traditional varieties (Pautasso et al. 2013), may also increase the overall resilience of resistance gene deployment if they deploy different types of resistance genes. Modularity can also be a weakness as individual decision makers in various nodes can prevent larger strategic initiatives from being implemented, and also limit exchange of varieties, priorities and knowledge, especially in cases where nontraditional actors and networks are particularly important.

A primary question when evaluating crop breeding networks is whether their structure results in resistance genes being developed and deployed for the optimal benefit to food security. The topology of crop breeding networks helps to determine whether this will be the case. For example, if there is a shift in directionality, such that the major international crop breeding groups change from being public groups with high out-degree (a high number of links leading outward from a node) to being private groups with high in-degree (a high number of links leading into a node), this may provide different challenges for optimal deployment of resistance genes (Fig. 3). For example, it may not be profitable to develop resistant varieties adapted to the needs of resource-poor farmers. However, private groups do not have their funding limited by availability of public funds and donor money, and if expected profitability can be demonstrated, more new cultivars may be produced. Alston et al. (2009) indicate that for agricultural research in general, much development benefiting resource-poor farmers has been a positive externality for research targeting wealthier farmers who constitute larger markets in wealthy countries. Analyses similar to those applied in the study of epidemic networks can also provide perspective on the likely success of crop breeding networks. One key aspect is the length of the lag between observed emergence of new pathogen types in the field, and network response to provide an elite cultivar with appropriate resistance. Epidemiology and risk assessment can contribute to shortening this lag time through regional and global monitoring and predictions of the breakdown of resistance genes, and of invasions of new pathogens, to provide more efficient feedback about epidemics to inform the priorities of crop breeders. Understanding the structure of epidemic networks, and how they change over time, can guide efficient monitoring (Sanatkar et al. 2015; Sutrave et al. 2012).

**Figure 3 / in Box 2.**
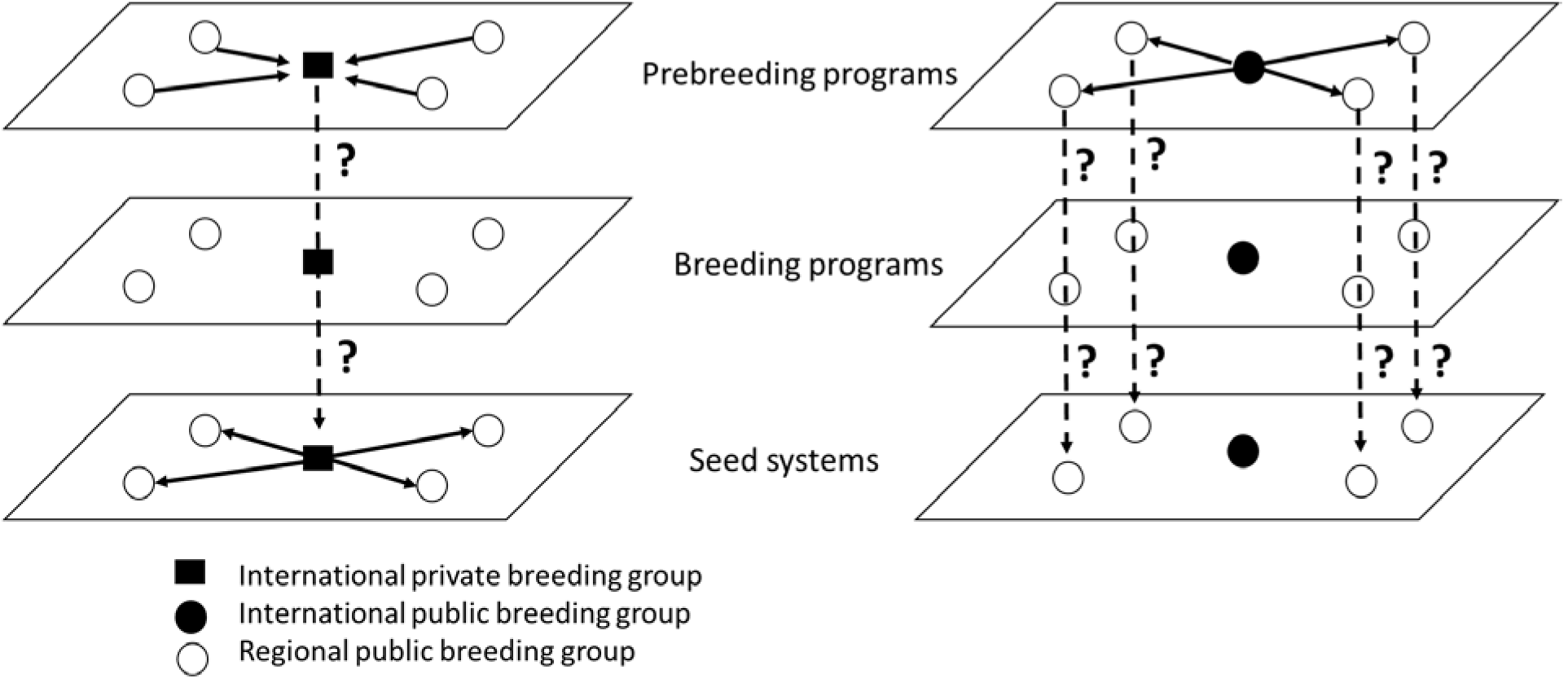
The **directionality** of links is a key feature of the multilayer networks that make resistance genes available for farmers to use. These schematic diagrams indicate two extreme cases and the potential limitations of such systems. In the first case (A), privatization of seed systems results in gene flow from prebreeding programs only into a private group. The question in this case is whether the private group will deploy the resistance genes in varieties that are readily available to farmers, or whether some genes that would have been useful, at least regionally, will not be available. In the second case (B), a centralized public breeding group provides gene flow in prebreeding materials throughout the system. The question in this case is whether the regional crop breeding groups will have enough crop breeders prepared to incorporate the resistance genes in regionally adapted varieties.

The coarse representation of the crop breeding networks here allows conclusions about the general structure of the networks and its implications for the deployment of resistance. More detailed analysis of specific crop networks will allow additional conclusions about whether the network structure is advantageous for particular regions and purposes. For example, from the standpoint of a particular region, how well does the network function to provide needed resistance genes, in enough variety to provide more lasting resistance? This can be evaluated in part by considering network features such as the node degree distribution, the frequency with which each potential node degree occurs. If a network is “scale-free” (Shaw and Pautasso 2014), there are nodes of high degree that help to maintain distribution through the network. CGIAR breeding groups have high out-degree but may have their capacity constrained by monetary resources, while private breeding groups may instead have high in-degree and low out-degree but access to more monetary resources. If a network is a “small world” network (Shaw and Pautasso 2014), there are links that cross different components of the network, keeping the number of steps between any two nodes low. Cut point analysis can identify nodes which, if removed, would substantially reduce the network coherence; in the current network, removal of CGIAR breeding groups would have a large effect to reduce coherence.

A key question is how readily new viable public and private breeding groups can arise to replace ones that might be lost from the system, and how readily new links between breeding groups can form to compensate for lost groups. The vulnerability of the ICARDA seed collections in Aleppo, Syria, is a striking example of the need for system redundancy (Bhattacharya 2016). Other traits of more detailed networks can be considered, such as controllability, the ability of a set of driver nodes to push the network toward a desired state (Liu et al. 2011). In the coarse perspective on networks presented here, it is clear that CGIAR breeding groups and large private breeding groups are key to controlling the state of the global crop breeding network. They are also likely to complement each other in some areas, and compete in others, and future research needs to clarify where and how in more detail. At a regional level, with more detailed information, there are likely secondary levels of key driver breeding groups, whose roles in network resilience would be useful topics for more detailed analysis.

The information networks associated with crop breeding networks, and innovation networks more broadly, are also a key system component (Garrett 2012, 2017; Poland 2015; Spielman et al. 2009). As gene editing technologies advance, the movement of physical materials may at some point no longer be necessary as information about DNA sequences alone may be enough for crop breeders to incorporate new forms of resistance. This development increases the importance of capacity in the different nodes, and without active policies, this may increase the competitiveness of private breeders and increase the challenges for breeders, in general, and especially for implementing joint strategies in developing countries.

The goal of system resilience has led to the development of a number of principles for improved systems (Table 1), where many concepts about resilience remain to be tested because the science of resilience is still in its early stages (Biggs et al. 2012; Donohue et al. 2016; Ostrom 2009; Scheffer et al. 2012). The scale of global crop breeding networks makes experimentation difficult. However, observational analyses of successes and failures in the history of resistance gene deployment, as a function of crop breeding network structure, could provide useful insights. Strengthening crop breeding programs in regions that are underserved by the current networks may improve the system-level outcomes of global crop breeding networks (Ribaut et al. 2010; Rivers et al. 2015), but the market incentives in better served regions may well increase the disparity in service levels. An important form of system-level research will also be to evaluate potential system interventions, to aid in prioritization when resources to invest in system improvement are limited and to provide incentives to provide public goods. Resistance gene deployment offers additional challenges for resource investment strategies, compared to genes such as those for drought tolerance, because deployment strategies used by some actors can reduce the utility of resistance genes for other actors in the system (Fig. 4).

**Table 1.** Principles for supporting system resilience, adapted fromBiggs et al. (2012), and potential application to global crop breeding networks. Some principles may apply to both the resistance genes, themselves, and the agents who disperse and deploy them.

| Principles from Biggs et al. (2012) | Principles applied to global crop breeding networks |
| --- | --- |
| 1. Diversity and redundancy | The benefits of deploying stacks of resistance genes that act independently, or strategic deployment of a range of resistance genes in mixtures, are well known. Mechanisms that result in different pools of resistance genes being deployed for different purposes may help to *create* useful forms of functional diversity for genes. A global crop breeding system with a critical mass of crop breeders, who represent a range of incentive structures and crop breeding strategies, may provide a form of insurance and make the system more likely to successfully adapt to the challenges of global change. For some systems, such as maize, crop breeding is highly privatized. For other systems, such as most tropical fruit crops, the very small number of crop breeders represents a system vulnerability. |
| 2. Management of connectivity | At the same time that the spread of resistance genes provides benefits for global resistance deployment, the spread of pathogens with crop germplasm, or crops postharvest (Hernandez Nopsa et al. 2015), is one obvious risk of connectivity among regions. Another potential risk of resistance gene dispersal is an increased probability of deployment in ways that decrease the useful life of genes (Figure 4). Information flows among crop breeders will <u>generally enhance decision-making and system success</u>. |
| 3. Management of feedbacks | Immediate feedbacks for crop breeders include the hectareage over which their new varieties are grown. Follow-on feedbacks in crop breeding networks include disease and pest responses to resistance gene deployment, and resulting profitability responses. Currently, potential regime shifts (Scheffer et al. 2012) for some crops may occur due to increased use of hybrids or herbicide-resistant varieties, with rapid feedbacks to privatize crop breeding systems. The resulting shift in incentives has the potential to push the system beyond a threshold, such that deployment and dispersal of resistance genes would slow as pesticide sales become more profitable. |
| 4. Fostering understanding of CAS; encouraging learning and experimentation | Understanding global crop breeding networks as systems may enhance the ability of planners and policy makers to strengthen the systems and prioritize resource investment. Conceptualizing crop breeding systems as part of broader systems providing ecosystem services may be useful, along with a longer-term perspective on gene stewardship that does not discount the future so strongly (Cheatham et al. 2009; Levin et al. 2012; Messier et al. 2014). |
| 5. Broadening participation | The active participation of a wider range of stakeholders could help to insure that the deployment of resistance genes meets the needs of farmers and of society more broadly. Effective information flows from farmers to crop breeders could also improve system responses to new stressors. |
| 6. Promoting polycentric governance systems | Having multiple “governing authorities” at multiple scales may improve crop breeding networks through effective regulation, in combination with market forces. Challenges include matching the scale of governance to the scale of potential problems, and making effective group decisions to handle trade-offs among the goals of different stakeholders (Biggs et al. 2012). |

**Figure 4 / in Box 3.**
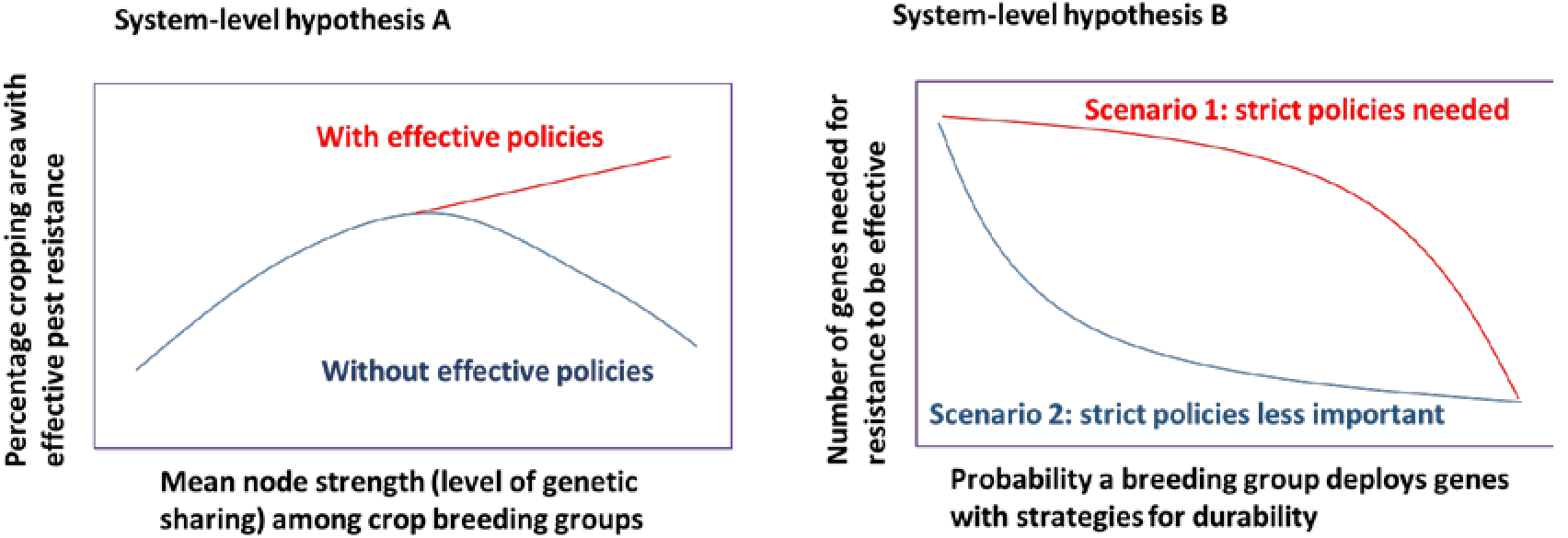
Hypotheses about resistance genes in cropping breeding networks. **(A)** Sharing of genetic resistance resources has practical risks and rewards, as well as the ethical implications of providing or withholding support for enhanced food security. As the level of sharing of genetic material among crop breeders increases, the percentage of cropping area in which effective resistance is deployed increases, up to a point. The increase in area with effective resistance is due to increased availability of resistance genes for breeding programs. The decrease in area with effective resistance at higher levels of sharing is due to the potential for breakdown in effectiveness of genes, as more breeders use the genes and thus the probability of deployment without a strategy to protect gene utility increases. Policies to support resistance gene stewardship can make the benefit of gene sharing continue through higher levels of sharing. **(B)** For different crop breeding systems, there may be different scenarios related to the importance of effective policies. In general, as the probability increases that any given breeding group uses effective strategies for gene deployment, the number of genes necessary for the system to be successful decreases. For some systems (Scenario 1), the decrease in the number of genes necessary may decline slowly, so that strict policies are needed even when most groups use effective strategies. For other systems, (Scenario 2), the decrease in the number of genes occurs with a small increase in the probability that breeding groups use effective strategies, so policies are less important. Use of QTLs rather than major genes for resistance may also make Scenario 2 more likely than Scenario 1.

In conclusion, we have illustrated, in low resolution, the crop breeding networks which determine the global pattern of resistance gene deployment for four major food crops. Finer resolution analyses of crop breeding networks will help to guide policies for better global and regional network structures, particularly in the context of agricultural development. A key research question is how policies can optimize the balance between public and private breeding groups, to provide the societal benefits of strategic resistance gene deployment in optimal landscape patterns for disease and pest management. There are exciting opportunities to draw on the growing field of network analysis as a source of input for strategies.

## Acknowledgements

We appreciate information about potato breeding networks from M. Bonierbale, about rice breeding networks from R. S. Zeigler, R. K. Singh, and F. Xie, about wheat breeding networks from G. Bai and S. Sehgal. This research was undertaken as part of, and funded by, the CGIAR Research Program on Roots, Tubers and Bananas (RTB) and supported by CGIAR Fund Donors http://www.cgiar.org/about-us/governing-2010-iune-2016/cgiar-fund/fund-donors-2/. We also appreciate support by the CGIAR Research Program for Roots, Tubers and Bananas, the CGIAR Research Program on Climate Change and Food Security (CCAFS), USDA APHIS grant 11- 8453-1483-CA, US NSF Grant EF-0525712 as part of the joint NSF-NIH Ecology of Infectious Disease program, and US NSF Grant DEB-0516046. Mention of trade names or commercial products in this publication is solely for the purpose of providing specific information and does not imply recommendation or endorsement by the U.S. Department of Agriculture. USDA is an equal opportunity provider and employer.

